# Glioma stem cells invasive phenotype at optimal stiffness is driven by MGAT5 dependent mechanosensing

**DOI:** 10.1101/2020.09.25.313783

**Authors:** Emilie Marhuenda, Christine Fabre, Cunjie Zhang, Martà Martin-Fernandez, Thomas Iskratsch, Ali Saleh, Luc Bauchet, Julien Cambedouzou, Jean-Philippe Hugnot, Hugues Duffau, James W. Dennis, David Cornu, Norbert Bakalara

## Abstract

Glioblastomas stem-like cells (GSCs) by invading the brain parenchyma escape resection and radiotherapy. GSC invasion is associated with altered N-glycosylation pattern of integrins and other transmembrane proteins resulting in changed mechanosensing but details are elusive. Because the tumour microenvironment has an increased stiffness we studied the interaction between matrix stiffness, N-glycosylation and GSC migration. To mimic the fibrillar microenvironments, we designed 3D-ex-polyacrylonitrile nanofibers scaffolds (NFS) with adjustable stiffnesses by loading multiwall carbon nanotubes (MWCNT). We found that migration of GSCs was maximum at 166 kPa. Migration rate was correlated with cell shape, expression of focal adhesion (FA), Epithelial to Mesenchymal Transition (EMT) proteins and (β1,6) branched N-glycan binding, galectin-3. Mutation of MGAT5 in GSC inhibited N-glycans (β1–6) branching, suppressed the stiffness dependence of FA and EMT protein expression as well as migration on 166kPa NFS; underpinning the role of multibranched N-glycans as a critical regulator of mechanotransduction by GSC.

**Significance Statement:** During pathological processes in which cell migration is involved, cells undergo important functional changes in protein glycosylation and are responsive to environmental mechanical modifications. We addressed the question of the glycosylation role in mechanotransduction regulation of glioma stem cells. We created a bio-inspired 3D nanofiber scaffold (NFS) loaded with multiwall carbon nanotubes to obtain NFS of adjustable stiffness in physiological and pathological ranges. We highlighted and described a mechanism of fine mechanotransduction leading to a nonlinear migration response regarding to 3D microenvironment stiffness values. We show the importance to develop mechano-pharmacology as new therapeutic target by demonstrating the relationship existing between environmental stiffness and multibranched N-glycans catalysed by the MGAT5 enzyme to optimize directed migration.

## Introduction

New concepts are emerging taking into account the role of glycocalyx and N-glycosylation in mechanosensing process due to cell-ECM interactions, but there is no report demonstrating the direct role of a specific glycosylation enzyme regulating mechanotransduction and optimal directed migration.

Coordination between FA dynamics and actin cytoskeleton is essential for cell migration (1). FAs are the adhesion nexus between cells and the ECM (2), which consists of a series of dynamic protein complexes interactions (3) highly modulated by stiffness and mediate mechanosensing. The molecular pathways for force transmission through the FA depend on direct interactions between the ECM and integrins (4) as well as adaptor proteins that directly connect integrins to the actin cytoskeleton, such as talin (5), indirect interactions between integrins and actin mediated by vinculin (6) and signalling pathways such as the Focal Adhesion Kinase (FAK) (7). Calpain2, an intracellular calcium-dependent cysteine protease, is involved in FA dynamics through modulation of talin and FAK structure and function (8) when bound to integrins (9).

Integrins have multiple site of N-glycosylation that are remodelled in the Golgi, modulating their affinity for galectins. The galectins bind to the N-acetyllactosamine (LacNAc) epitopes, with affinities that increase with N-glycan branching, catalysed by the Golgi N-acetylglucosaminyltransferase (MGAT) pathway comprising MGAT1, MGAT2, MGAT4a,b and MGAT5. The highly branched N-glycans catalysed by MGAT5 harbour the higher affinities for galectins (10, 11). The resulting galectin lattice displays rapid exchange of binding partners, thereby acting as an intermediary between free diffusion of glycoproteins, and more stable complexes notably the integrin-ECM focal adhesions (9, 12).

ECM stiffness is reported to increase from low grade glioma to GBM (13), whereby tissue stiffness has been suggested to modulate their migration capacity (14). These glioma-induced changes in ECM stiffness are accompanied by changes of the glioma cell morphology and nuclear volume (15). Clinical observations by Scherer (1938) (16), demonstrated that glioma cells preferentially migrate in fibrous areas (14). The natural ECM fibrillar architecture is critical for 3D mechanosignalling events such as FA formation (17) and fibre alignment has therefore been reported to greatly influence migration. However, little is known about stiffness impact of fibrillar 3D environment on migration.

We initially developed a NFS made of 3D electrospun ex-polyacrylonitrile which supported GSC migration *in vitro* reflecting the behaviour of glioblastomas *in vivo* (18). Beyond specifically mimicking fibrous structures the NFS includes a 3D topology around the cells that reproduce heterogeneity, directionality, surface chemistries relevant to understand the behaviour of GSCs (18). Substratum fibril composition and MGAT5 expression individually have been shown to influence cell adhesion and migration with non-monotonic dynamics (9, 12). In this paper we report the mechanosensing interaction existing between substratum stiffness and MGAT5 activity by profiling wild-type (WT GSC) and MGAT5 knock-out GSCs (MGAT5 KO GSC) seeded onto NFSs made with stiffnesses from 3 to 1260kPa, values in the range reported for human healthy tissues and gliomas (19). The results reveal a requirement for both MGAT5 branched N-glycans and an optimum substratum stiffness for mechanotransduction of signalling and GSC invasiveness.

## Results

### NFSs of different stiffness are generated with the addition of different amount of MWCNTs

Four grades of NFSs were prepared by incorporating different amounts of MWCNTs. These nanotubes present a very high intrinsic Young’s modulus and are consequently used to increase the Young’s modulus of composite fibres (20). The different NFSs respectively featured 0, 0.0015, 0.00635 and 0.05 % w/w of MWCNTs in order to recapitulate a comparable cellular environment. To confirm the stiffness, atomic force microscope measurements were done in a liquid medium checking that the contact point was well applied on the surface of the fibre. For each grade, at least 5 areas were analysed (Fig S1). These results yield average Young’s moduli of 3 ± 2 kPa, 166 ± 29 kPa, 542 ± 90 kPa and 1260 ± 430 kPa, respectively, indicating an increase in stiffness as a function of MWCNT content (Fig 1A).

**Figure 1:**
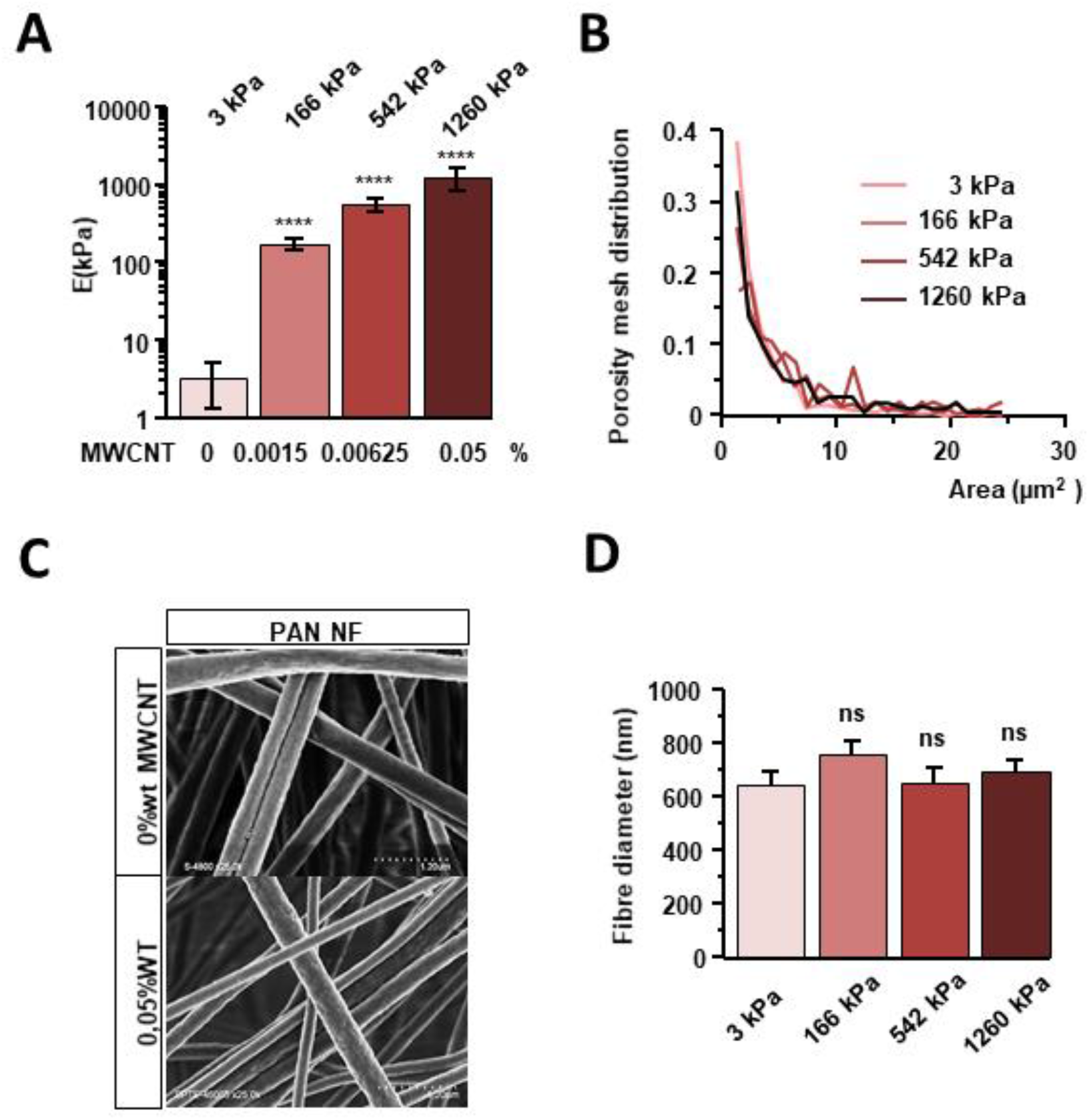
Characterization of the NFS with MWCNTs. (A) Young’s modulus increases as a function of MWCNT content. Bars represent the average of the Gaussian peaks fitted to the Young’s modulus value distributions of individual force maps (± SD). The number of analysed force maps is at least 5. The stiffness of the four samples are significantly different from one another (p < 0.01, one-way ANOVA). (B) Porosity distribution measured by confocal microscopy and 3D reconstitution. (C) Scanning electron microscopy images of fibres surface. (Scale 1.20μm) (D) Fibre diameter distribution (± SEM). (* p<0,05; ** p<0,01; *** p<0,005; **** p<0,001) Statistical significance was determined using one-way ANOVA with post hoc Tukey’s Honest Significant Difference test for multiple comparisons.

The electrospinning technic allowed us to produce a NFS creating a biomimicking, confined environment for embedded cells as previously described (18). Its pore area distribution ranges between 0.5 μm^2^ and 7 μm^2^, with constant fibre morphology and diameter, independent of MWCNT content (Fig 1B, C and D).

### 166 kPa NFSs trigger optimal GSC motility

NS of identical sizes were plated on NFSs of different stiffnesses and were allowed to migrate in differentiation medium for 5 days (Fig 2A and B). GSCs grow at the same rate on NFS of different stiffnesses (Fig 2C). GSCs migrate on NFS in a collective mode, irrespective of the stiffness of the fibres (Fig 2B and D). However, on the 166 kPa nanofibers a large number of GSCs migrated out of the NS, while migration was minimal on other nanofibers stiffnesses, and the migration area on 166 kPa NFS was ~4 times larger than on NFSs of other stiffnesses (Fig 2A, and E). We found similar results both in proliferation medium and differentiation medium. Proliferation medium is serum-free and supplemented with β-FGF and EGF which allows propagation of multipotent, self-renewing tumour-spheres (NS) (21). On tissue culture, the cells remain largely stationary as NS in this medium, but on 166 kPa nanofibers the glioma cells migrated (Fig 2A and F). The optimal stiffness of 166 kPa is sufficient to trigger migration, something which is usually only achieved by the addition of serum to the medium. Cytotoxicity or changes in proliferation were not observed on NFSs of different stiffnesses.

**Figure 2:**
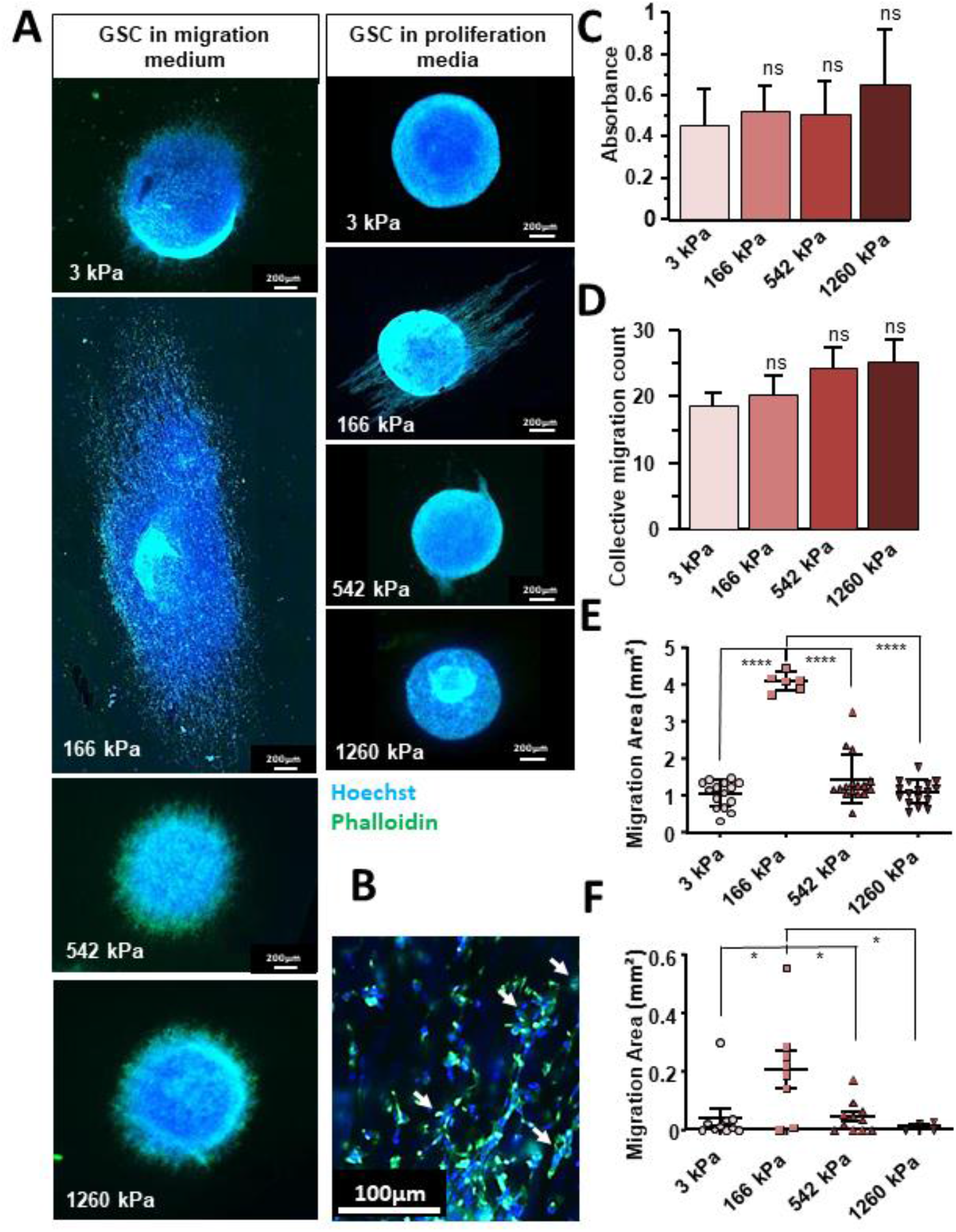
Optimal migration of GSCs on 166 kPa NFS. Migration was estimated by subtracting the area of the initial NS to the total area occupied by the NS and cells having migrated away from it within 5 days. (A, B) F-Actin was stained with the Phalloïdin (Green) and the nucleus stained with the Hoechst 33342 (Blue). (A) Left panel: in migration medium (Scale bar: 200μm). Right panel: in proliferation medium (Scale bar: 200μm). (B) A view of cells migrating on the 166 kPa NFS at higher magnification. Collective migration were considerate for each aggregates composed at least by tens cells tightly associated (18). Arrows point at a few examples of collective migration. (Scale bar: 100μm) (C) 3000 dissociated GSCs were seeded on NFSs of different stiffnesses and proliferation was estimated with the MTT assay after 5 days of culture. No significant differences between conditions could be discerned (n=3) (D) Collective migration quantification Migration areas in differentiation medium as a function of NFS stiffness showing maximum migration at 166 kPa (n=3 independent experiments, with at least 6 NS per condition) (E) Migration areas in migration medium as a function of NFS stiffness showing appreciable migration only at 166kPa (n=3 with at least 6 GSCs NS per condition). (F) Migration areas in proliferation medium as a function of NF stiffness showing appreciable migration only at 166kPa (n=3 with at least 4 GSCs NS per condition). Values of the figure correspond to the mean ± SEM and statistical significance was determined using one-way ANOVA, (* p<0,05; ** p<0,01; *** p<0,005; **** p<0,001).

### WT GSC morphology depends on NFS stiffness

Directionality of migration is reflected a bipolar spindle-shaped cell morphology driven by cytoskeleton rearrangement (15, 22, 23). Cell morphology was examined by plating dissociated GSCs on in 2D NFSs (Fig 3A), and after 5 days, we measured the cell length (Fig 3B), width (Fig 3C) and calculated width/length ratios (Fig 3D). GSCs plated on NFSs were always thinner than GSC plated in 2D. The length of the GSCs cultivated on NFSs of 542 and 1260 kPa were shorter than cells on NFSs of 3 kPa and 166 kPa. The minimum width was observed for cells on NFS of 166 kPa (Fig 3A and C). The width/length ratio indicates that the maximum morphological plasticity is reached for a stiffness of 166 kPa (Fig 3D). Cancer cell motility is associated with FA dynamics, and EMT phenotype (14, 24). We therefore investigated the expression of proteins implicated in EMT and/or cytoskeleton organisation next.

**Figure 3:**
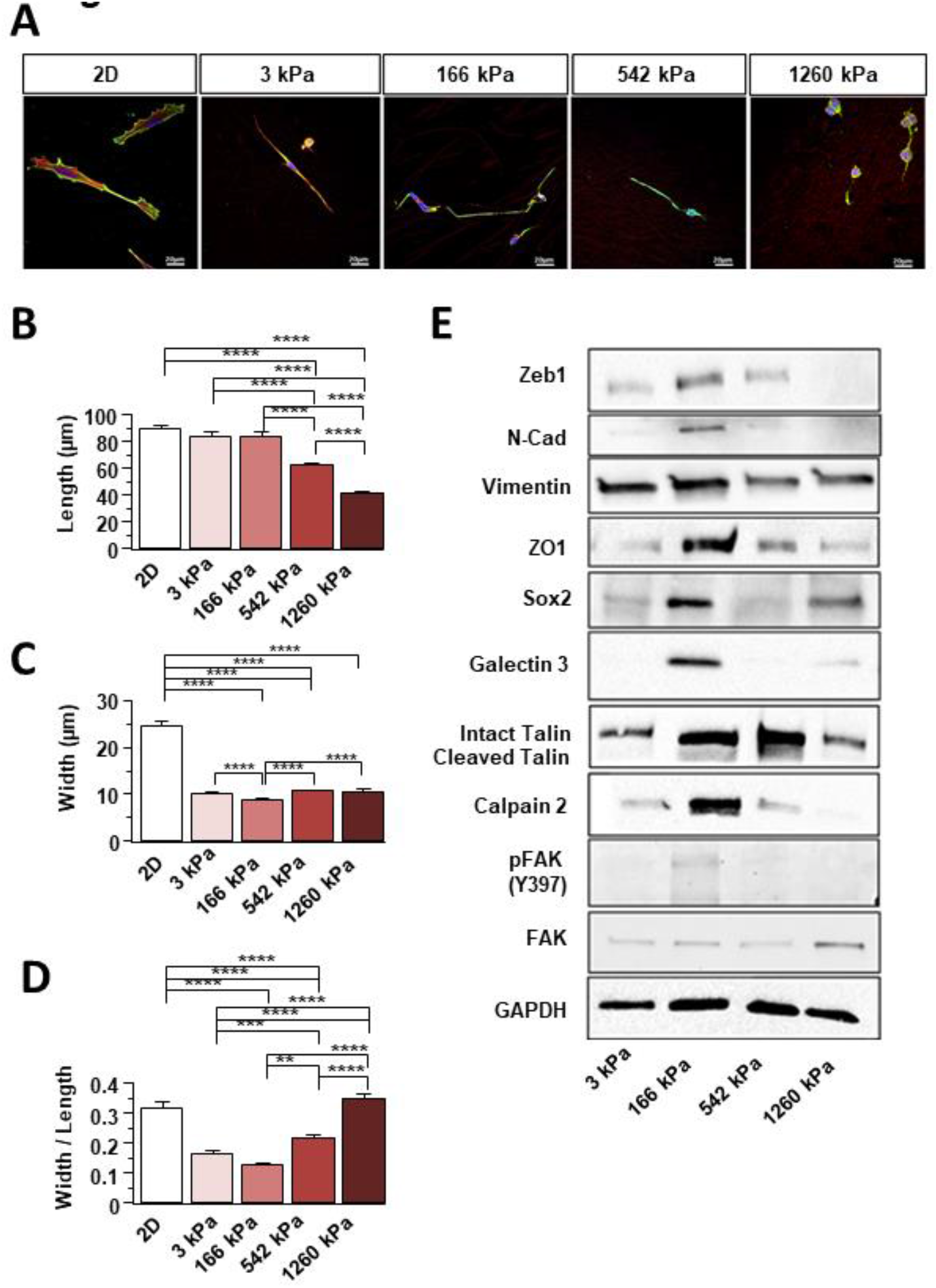
Optimal stiffness increases EMT and FA maturation in GSCs. (A) GSCs were cultured on 2D and NFSs of different stiffnesses. Labelling: vinculin green, actin red (phalloidin) and DNA blue (Hoechst 33342). (B) Length, (C) Width and (D) aspect ratio of GSCs grown fibres of different stiffnesses were determined. Measurements were taken regarding to nucleus location. (n=3 with at least 172 cells by condition) Values of the figure correspond to the mean ± SEM and statistical significance was determined using one-way ANOVA (* p<0,05; ** p<0,01; *** p<0,005; **** p<0,001). (E) Western Blot membranes showing EMT and focal adhesion protein expression in GSCs grown on NFSs with different stiffnesses. (n=3 to n=7 for each protein tested) Associated quantifications in Supp. Fig. 2.

### Expression of EMT and adhesome proteins is maximal on the 166 kPa NFS

The expression of N-CAD, ZEB1 and SOX2 was higher in cells grown on 166 kPa NFS than in cells cultured on the other NFSs (Fig 3E and Fig S2). This agrees well with the higher motility GSCs observed on 166 kPa NFS. Although, vimentin expression does not vary significantly with fibre stiffness (Fig 3E and Fig S2), the tight junction protein ZO1 was selectively expressed in cells grown on 166 kPa NFS (Fig 3E and Fig S2). These data suggest that cell-cell junctions and their turnover may play a role in efficient collective migration at the optimal stiffness of 166 kPa.

FA maturation and signalling through integrin-ECM interactions, as well as FAK signalling modulate cell migration (12, 25). FAK phosphorylation on tyrosine 397 (Y397) is associated with integrin mechanosensing as well as with integrin-mediated focal adhesion maturation and turnover (26). The pFAK/FAK ratio was elevated in cells grown on 166 kPa NFS compared with the NFSs of different stiffness (Fig 3E and Fig S2), consistent with the relative migration rates (Fig 2). The levels of calpain 2, talin and its cleaved form are also highest in cells cultured on 166 kPa NFS. These proteins are recruited to FA where calpain 2 promotes the recycling of these structures and contribute to migration (8, 26). Integrins are modified with β1,6GlcNAc branched tetra-antennary N-glycans are bound by Galectin-3 in a transient interaction that promotes FA remodelling depending also on stiffness of the ECM (9, 12, 27). Galectin 3 levels are increased in cells grown on 166kPa NFS (Fig 3E and Fig S2). Both MGAT5 and Galectin-3 are upregulated in transformed cells, and causal associated with invasion and metastasis (12). Our results suggest that their interaction may be required for mechanosensing of ECM stiffness and cell migration. To explore this hypothesis, MGAT5 was deleted in GSC clones using CRISPR-Cas9 (28) and the absence of the enzyme checked by western blotting (Fig 4A).

**Figure 4:**
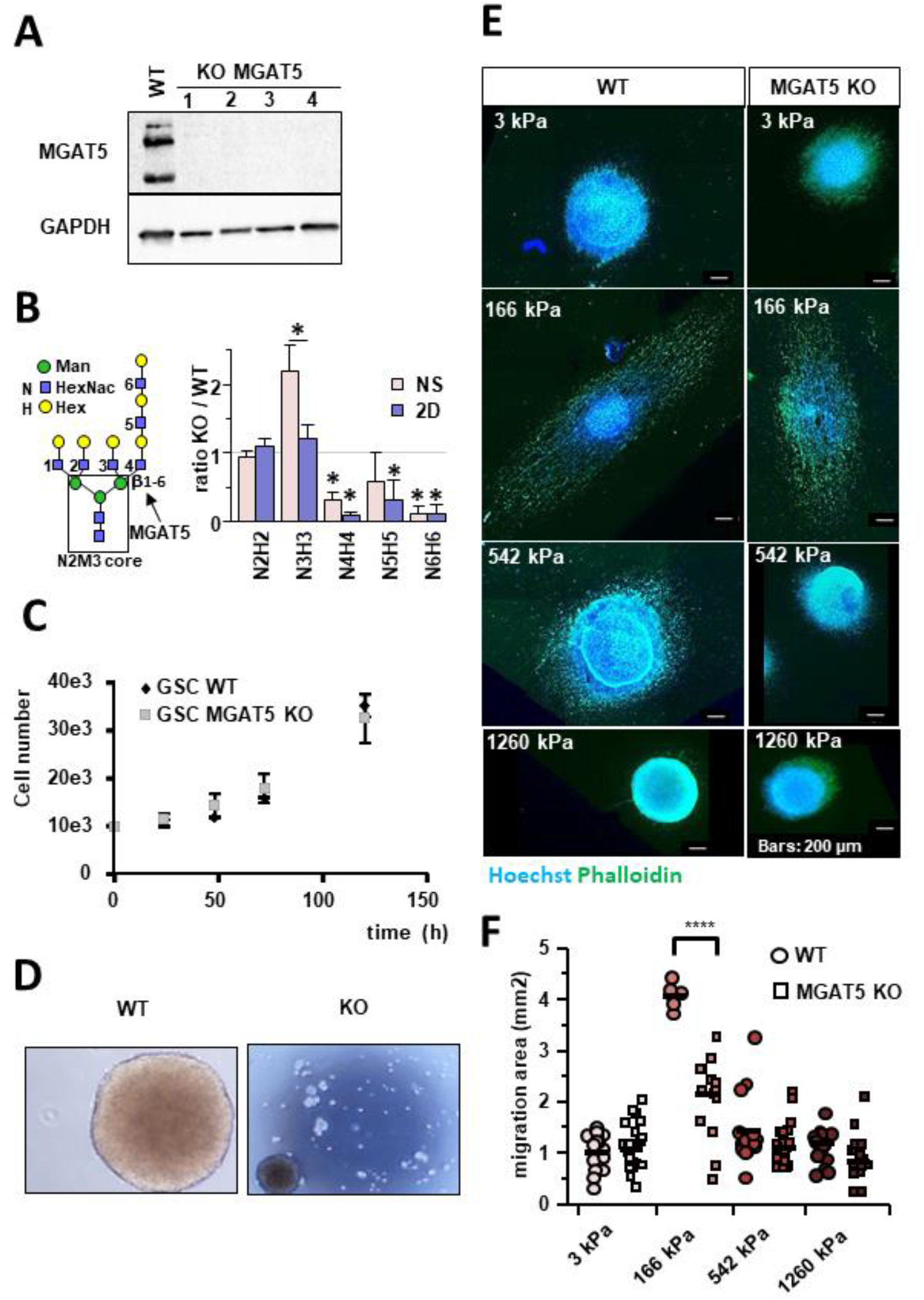
KO of MGAT5 decreases GSC migration at 166 kPa. (A) Western blot of MGAT5 expression in WT and 4 MGAT5 KO clones. (n=3) (B) LC-MS/MS data for total bi- (N2H2), tri- (N3H3) and tetra- (N4H4) antennary N-glycans expressed as a ratio of MGAT5 KO / WT reveals the loss of N4H4 and addition of polylactosamine >N4H4; (* p<0.05). Our analysis does not distinguish lactosamine isomer, which are therefore represented as minor structures in structures with >N3H3 units. The N-glycans with masses >N3H3 in KO cells, are most likely N2H2 and N3H3 with polylactosamine extensions with identical masses to N4H4. (n=3) (C) Proliferation of WT and MGAT KO GSCs does not differ (n=5). (D) Compared to WT GSC, NS formation is inhibited in MGAT5 KO GSCs. (E) Migration of WT and MGAT KO GSCs on fibres of various stiffnesses. F-Actin was stained with Phalloïdin (Green) and the nucleus stain with Hoechst 33342 (Blue) (F) Quantification of migration area in differentiation medium comparing WT and MGAT5 KO GSCs showing a significant reduction of migration of KOs at 166 kPa (n=3 with at least 6 GSCs NS per condition). Values of the figure correspond to the mean ± SEM and statistical significance was determined using one-way ANOVA (* p<0,05; ** p<0,01; *** p<0,005; **** p<0,001).

### MGAT5 knock out decreases migration on the 166 kPa NFS

Glycome analysis showed that β1,6GlcNAc branched tetra-antennary N-glycans were absent in the KO cells with respect to WT, while the relative expression of triantennary N-glycans was increased (Fig 4B). The major fucosylated N2M3+N4H4F isoform group 2 is much less abundant in MGAT5 KO compared to WT (Fig S3). The remaining N2M3+N4H4F structures observed in the MGAT5 KO GSCs (Fig 4B) were most likely isomers equivalent in mass, notably bi- and tri- antennary with polylactosamine (Fig S3).

The MGAT5 KO GSCs grow at the same rate as wild type GSCs (Fig 4C). However, in proliferation medium they formed multiple small and weakly adhering spheres rather than a single NS as seen for WT, suggesting that cell-to-cell adhesion is compromised (Fig 4D). We previously reported a similar effect on NS formation with the MGAT5 inhibitor, phostine PST 3.1a on GSCs (29). In agreement with this, we also observed a disaggregation of the MGAT5 KO GSCs NS and dispersion of the cells on NFSs, when grown in proliferation medium (Fig S4).

In differentiation medium, which promoted migration of WT GSC, migration of MGAT5 KO GSCs was significantly reduced on the 166kPa NFS only (~ 2 times) (Fig 4E and F), highlighting the implication of MGAT5 activity in optimal rigidity sensing. Whereas cell dimensions of WT GSCs show a clear dependence on matrix stiffness, KO cells change shape much less as a function of stiffness (Fig 5 A-D).

**Figure 5:**
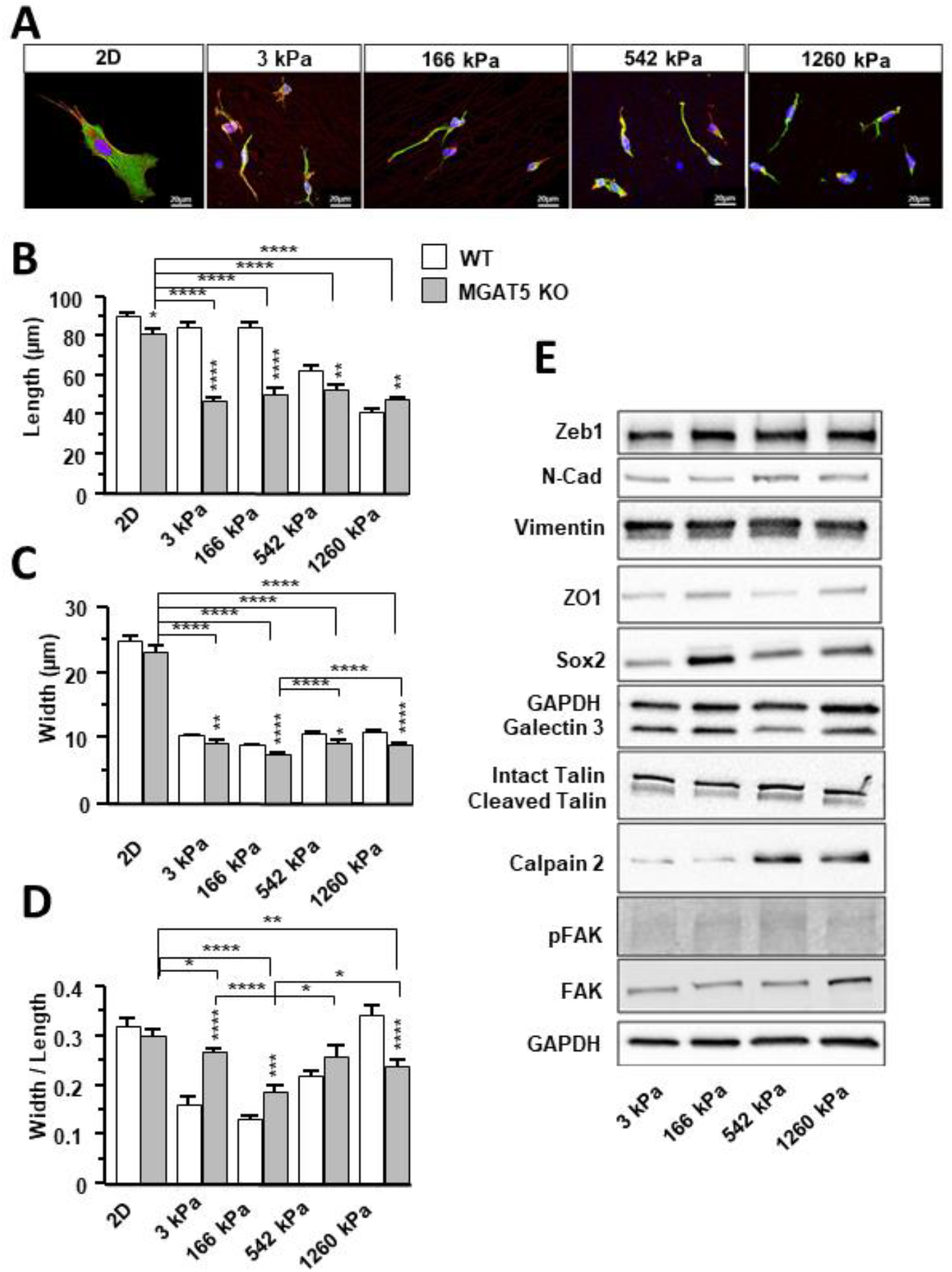
MGAT5 KO modifies the expression of EMT and adhesome proteins. (A) MGAT5 KO GSCs were cultured on 2D and NFSs of different stiffnesses. Labelling: vinculin green, actin red (phalloidin) and DNA blue (Hoechst 33342). (B) Length, (C) Width and (D) aspect ratio of MGAT5 KO GSCs grown fibres of different stiffnesses were determined and compared to WT. Asterisks indicate significant differences between KO and WT. (n=3 with at least 86 cells by condition) (E) Western Blot of EMT and adhesome protein expression by MGAT5 KO GSCs (n=3 to n=4 for each protein tested). Associated quantifications in Supp. Fig. 4. Values of the figure correspond to the mean ± SEM and statistical significance was determined using one-way ANOVA (* p<0,05; ** p<0,01; *** p<0,005; **** p<0,001).

The length and width of the MGAT5 KO GSCs did not vary according to the stiffness of the NFSs (Fig 5B and C). Width/length ratios for MGAT5 KO GSCs compared to those of WT GSCs (Fig 5 D), revealing a reduction in morphological plasticity on 166kPa NFS.

The robust increases in ZEB1, N-CAD, SOX2, ZO1, Galectin 3, and Talin, Calpain2 and pFAK/FAK observed for WT GSCs on 166 kPa NFS were not observed in MGAT5 KO GCS with the exception of SOX2 (Fig 5E, 3E and Fig S2). Thus FA maturation and signalling in MGAT5 KO cells was inhibited on 166 kPa NFS and comparable to other NFSs stiffnesses. Thus MGAT5 N-glycan branching is critically involved in glioblastoma mechanosensing and regulation of stiffness dependent cell morphology and motility (Fig 6).

**Figure 6:**
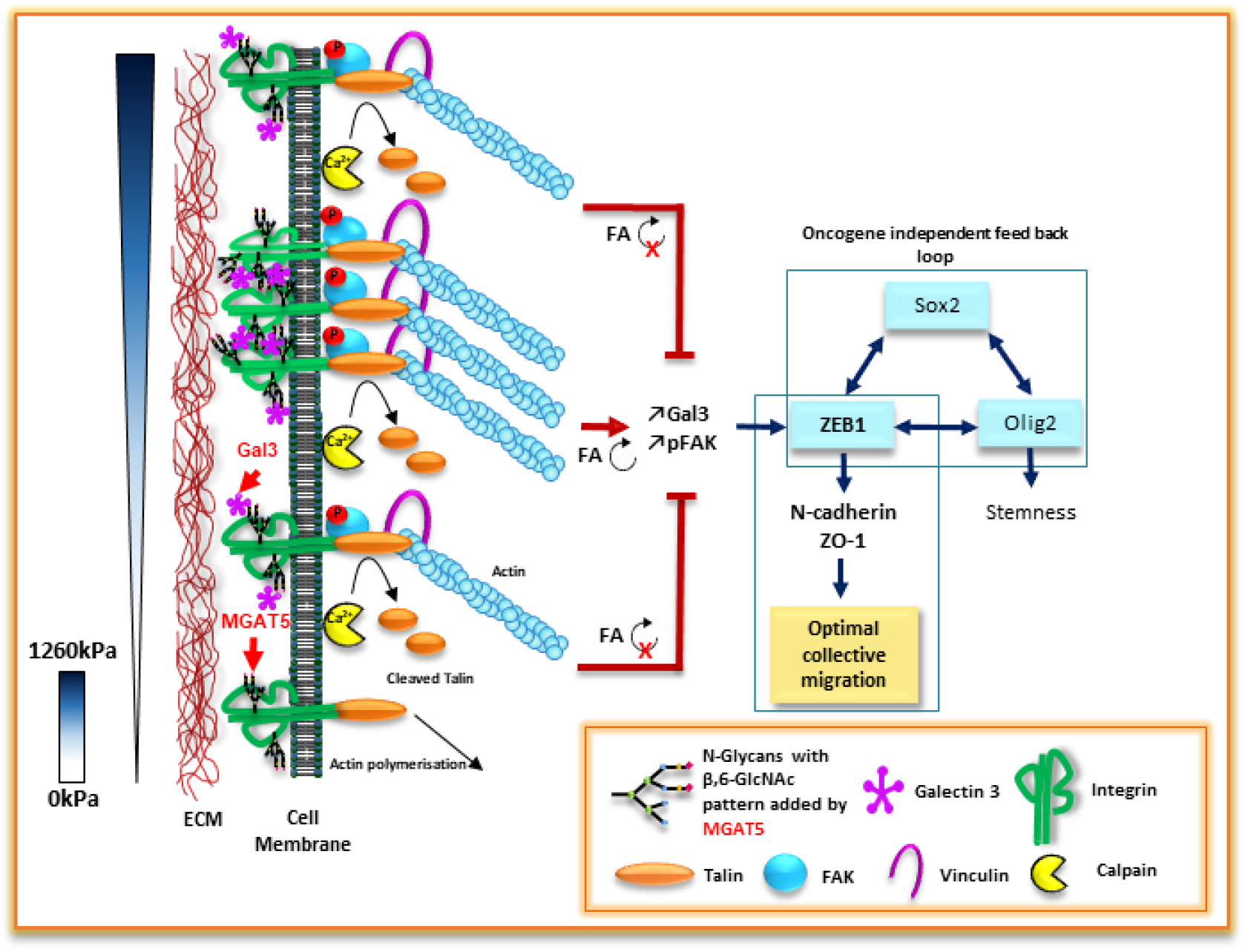
MGAT5 involvement in mechanotransduction and cell migration rate control. 166 kPa is the optimal stiffness for integrin clustering controlled by MGAT5 glycosylation activity, focal adhesions (FA) turn-over and maturation (↻) and galectin 3 (Gal3) up-regulation. The resulting augmentation of FAK signalling increases expression of the mesenchymal marker ZEB-1 with subsequent increases of N-CAD and ZO-1 expressions leading to increased cell collective migration rate. ZEB-1 appears as a hub integrating signalling information from the SOX2 transcription circuit, independent of the RTKs (50) and the MGAT5 restricted mechanotransduction depending on ECM stiffness (S).

## Discussion

Tumor cells undergo functionally important changes in protein glycosylation (30, 31), notably, increased N-glycan branching and extension with polylactosamine, the preferred ligands for galectins and regulation of cell surface receptors (10, 11). The galectin interaction with branched N-glycans on integrins is associated with turnover and maturation of the FA, fibronectin fibrillogensis and actin microfilament remodelling (12, 27). Knockdown of MGAT5 in cancer cells has been shown to reduce N-glycan branched on N-CAD and turnover of cell-cell adhesions, as well as cell migration (32). Removal of three sites with branched N-glycans from N-CAD did not alter surface expression but did reduced cancer cell migration and invasion. Here we examined the role of MGAT5 in the sensing of substratum stiffness by GSC, through FA formation and signalling. GSCs displayed an optimal migration on 166kPa NFS relative to higher or lower stiffness. Moreover, increases in galectin-3, talin, calpain2 and the pFAK/FAK ratio in WT GSC the 166Pa NFS indicate optimal FA signalling and turnover consistent with the dynamics of galectin-lattice model in cell migration (9, 12). We observed that a stiffness of 166 kPa is sensed as optimal for migration even when the chemical signals given by the proliferation culture medium constrain them to NS phenotype. This critical value of 166 kPa illustrates the physical crosstalk existing between cell applied forces and stiffnesses of the ECM to reach a maximum migration speed. Less or more stiffness do not appear to allow FA maturation and maximal cell migration. The increased amount of talin and cleaved talin observed using 166 kPa NFS reflects the strengthening of the link between the integrin and the actin cytoskeleton in FAs (33, 34) and illustrates a role for talin as a tension sensor (35). In addition, Calpain which expression increases in 166 kPa NFS is known to cleave talin and FAK, thereby regulating both adhesome dynamics (8) and rigidity sensing (33). In conclusion, measurement of the ECM stiffness and MGAT5-dependent N-glycan branching may be a better indicator of cancer cell invasiveness than either alone.

β1,6GlcNAc branched N-glycans with polylactosamine extensions have higher affinities for galectin-3 than less branched structures (11). We also noted that unlike NS, the relative expression of N3 branched glycans does not increase for 2D MGAT5 KO GSCs. Because formation and disassembly of integrin-mediated adhesions are regulated temporally and spatially during migration (36) we hypothesize the relative amounts of tri- and tetra-antennary N-glycans control migration.

Interestingly, loss of MGAT5 expression severely reduced GSC migration speed on 166 kPa NFS only and enhanced cell detachment from the NS generating cell dissemination. Cell detachment and dissemination of MGAT5 KO GSCs is also observed and increased when cells seeded on NFS of different stiffnesses grow in proliferation medium. In contrast to what was observed in WT GSCs, Galectin3 expression profile and pFAK/FAK ratio in MGAT5 KO GSCs, didn’t vary according to NFS rigidity. These results are in agreement with our previous data showing that inhibition of MGAT5 enzymatic activity by the phostine compound PST3.1a causes cell detachment and dysregulation of actin assembly subsequent to a reduction of FAK phosphorylation thereby causing reduced migration of Gli4 GSCs (29). These data highlight the key role played by MGAT5 in regulation GSCs migration speed via FAK signalling. Cell migration is also a remarkable example of the close link existing between integrin signalling, assembly of the cytoskeleton and cell morphology (37). In MGAT5 KO GSCs compared to WT GSCs, we observed a decrease in GSCs length. This morphological difference is associated with a decrease of calpain2 and talin expression on 166kPa NFS. Those results suggest a loss of FA maturation mediated by stiffness on MGAT5 KO GSCs. They also underline the impact of the NFS stiffness on galectin 3 expression modulation mediated by FA maturation and turn-over associated with MGAT5 expression. In conclusion these data are consistent with the reported observations that cell migration is the result of cell attachment and detachment regulated by the FA turnover in which FAK also plays a central role (38).

ZEB1 expression has been reported to play a central role in regulating GSCs migration (39). The expression of ZEB1 did not vary with rigidity in MGAT5 KO GSCs compared to WT GSCs). In addition, the absence of ZEB1 expression modulation according to the different NFS rigidities highlights that MGAT5-mediated glycosylation is essential for mechanical environment sensing supported by a differentially regulated expression of ZEB1. ZEB1 was reported to enhance the expression of the mesenchymal marker N-CAD (40). We observed that enhanced expression of ZEB1 on 166 kPa NFS in WT is associated with an increase of N-CAD expression. In MGAT5 KO GSCs the loss of the ZEB1 expression modulation according to the NFS stiffness is accompanied by a loss in the differential expression of N-CAD. These results confirm that MGAT5-mediated glycosylation promotes the sensing of the mechanical properties of the microenvironment through FA maturation and promotes EMT at 166 kPa. Nevertheless, in MGAT5 KO GSC grown on 1260 kPa NFS, we observed both an increase in Calpain2 and N-CAD expressions. The latter data indicates that in absence of MGAT5 expression, higher stiffness is required for membrane deformation in order to induce FA turnover and EMT-like phenotype.

In addition to the cell-ECM interactions and signaling, cell-cell interactions, play a key role in collective cell migration (41) and N-CAD is reported to promote cancer cell migration (42). In contrast to WT GSCs, ZO1 expression was down regulated in MGAT5 KO GSCs in NFS of the optimal stiffness of 166 kPa. Therefore we suggest that a deficient galectin lattice, lead to uneffective transmission of mechanical information orchestrated by FAs and cell-cell junctions unabling optimal migration and stiffness dependant EMT.

Beside the effects on cell migration and mesenchymal in MGAT5 KO GSCS the expression of SOX2 remains inducible according to the rigidity of the support as in WT GSCs. This result suggests that part of the stemness character is conserved independently of MGAT5 expression. In addition, we show that the action of SOX2 to reach a maximal velocity on NFS of stiffness of 166 kPa is restricted to MGAT5 glycosylation activity and to ZEB1 expression modulation according to stiffnesses.

In conclusion, the absence of MGAT5 leads to a decrease in migration speed, EMT-like processes and FA turn-over and maturation through the inhibition of mechanotransduction. We propose that the mechanisms involving MGAT5 in EMT processes is mediated by fine sensing of the stiffness leading to ZEB1 expression modulation, and regulate key oncogenic functions (43), making MGAT5 as a serious target to treat cancer.

## Materials and methods

### 3D nanofibre matrix

Polyacrylonitrile nanofibers were produced by electrospinning using a solution of 10 % W/W Polyacrylonitrile (Sigma Aldrich) dissolved in dimethylformamide (DMF, Sigma Aldrich). The electrospinning set-up (IME Medical Electrospinning, NL) was equipped with a rotating drum module located at a distance of 15 cm from the needle. In a classical experiment, a voltage of 20 kV was applied and the rotating speed of the drum was set at 2000 rpm. To tune NFS mechanical properties, multiwalled carbon nanotubes (MWCNTs, Nanocyl, 95% purity) were added in the Polyacrylonitrile solution and stabilized by adding Triton x100 (Sigma Aldrich) at a weight ratio of 1:50 of MWCNT/Triton. After electrospinning, the NFSs were cross-linked by heat treatment at 250°C (4 °C/min) during 2 hours under air. Pieces of the NFSs were cut and sterilized using classical autoclave treatment before further biological use (18).

### Atomic Force Microscopy (AFM)

Atomic force microscopy measurements were done using an Asylum MFP-3D head coupled to a Molecular Force Probe 3D controller (Asylum Research, Santa Barbara, CA, USA). Triangular silicon nitride cantilevers (MLCT, Veeco) with a nominal spring constant of 10 pN/nm and half-opening angle of 35° were used. The probe has a nominal length of 310 μm, width of 20 nm, and resonance frequency of 7 kHz. Prior to each experiment, the cantilever spring constant was determined in liquid using the thermal noise method available within the MFP-3D software. Samples were glued to a Petri dish by means of carbon conductive double-faced adhesive tape, in order to minimize electrostatic interactions between the atomic force microscope tip and the sample, and were covered with 2ml of deionized water. After testing a range of loading forces on the sample surface, the measurements were performed in liquid and at room temperature with a maximum loading force of ~300 pN corresponding to a maximal indentation depth of 120 nm. Higher loading force values (up to 1 nN) led to a stiffness overestimation due to a possible influence of the substrate.

The elastic deformation was obtained from the force curves as a function of the loading force applied by the tip. Young’s modulus (E) was calculated for each force from the approaching part of the force curves as the recorded force curves exhibited hysteresis. It shows the viscoelastic behaviour of the sample (44), following a modified Hertz model (45), based on the work of Sneddon and further developed for different atomic force microscope tip shapes as described elsewhere (46). A constant approach velocity of 6 μm/s was chosen, meaning a piezo-extension rate of 3 Hz to minimize hydrodynamic and viscoelastic artefacts (47). The Poisson’s ratio of the cells was assumed to be 0.5, as suggested for incompressible materials (48). The analysis of one image for each type of NFS is presented in supplementary figure 1.

### Cell culture

GBM cells isolation and primary cell culture were realized using the classical non-adherent neurospheres (NS) protocol adapted by Guichet *et al* (49). GBM cells were cultured in two different conditions in DMEM/F12 medium supplemented with glucose, glutamine, insulin, N2 and ciprofloxacin. In the “proliferation” non-adherent condition, the culture flasks were pre-coated with poly-2-hydroxyethyl methacrylate (poly-HEME, Sigma) and the medium was also supplemented with Epidermal Growth Factor (EGF), Fibroblast Growth Factor (FGF), gentamycin, heparin, fungizone, fungin and B27. In this condition, GBM cells growing as NS is reminiscent of neural stem cells *in vitro*, express neural progenitors and stem cells markers (nestin, OLIG2, SOX2 etc.), self-renew and propagate tumours in immunocompromised animals. NS could be dissociated and reseeded. In the “differentiation” condition, also called “migration medium”, the DMEM/F12 medium was supplemented with foetal bovine serum (0.5%), fungizone and B27. For this latter condition, GBM cells dissociated or as NS were cultured in adherence on 2D (planar surfaces). These cells are called adherent cells. Also GBM cells dissociated or as NS are cultivated on NFs of various stiffnesses. The GSCs remained in culture during 5 days at 37°C / 5%CO2.

To obtain NS with the same size, we used Corning^®^ 96 well round bottom ultra-low attachment microplates coated with a covalently bond hydrogel (Corning 7007). Dissociated GBM NS cultured in proliferation condition, were seeded at 7500 cells per well and remained in culture during 6 days until formation of single NS after sedimentation.

### MGAT5 knock-out

MGAT5 KO GSCs were made using the CRISPR Cas 9 technique (28). GSC transfection was carried out by electroporation with 1μg of CRISPR Cas 9 plasmid (Santa Cruz) per one million cells in Amaxa mouse neural stem cell Nucleofector. Transfected plasmid contained GFP for Fluorescence-activated cell sorting and to deposit one cell per well in Corning^®^ 96 wells round bottom ultra-low attachment microplates. After proliferation and the formation of a single neurosphere, KO efficiency was checked by western blot.

### Quantification of migration

NSs were deposited on NFSs of different stiffnesses in differentiation/migration medium and left to migrate for 5 days (18). Migration quantifications were done using ZEN 2012 software in counting number of cells, using DAPI staining. Migration capacity was quantified by measuring an area of migration. To measure migration areas, we subtracted the area of the NS containing non-migrating cells from the total area where cells were detected. Regarding collective migration, Gli4 cells were considerate as migrate collectively by forming aggregates composed at least of tens of tightly associated cells (18).

### Total cellular N-glycan profiling

Extracted membrane protein (30 μg) was suspended in 30 μL of 0.25% RapiGest SF, 50 mM ammonium bicarbonate, 5 mM DTT, heated at 85oC for 3 min, then mixed with 1 μL of PNGase F, 0.7 μL of sialidase, and 20 μL of 50 mM ammonium bicarbonate, and incubated at 42oC for 2 h followed by 37oC overnight. Released N-glycans were extracted with 4-5 volumes of 100% ethanol at −80° C for 2 hours. The supernatant containing released N-glycans was speed vacuumed to dry.

Pipet tips packed with 10 mg porous graphitized carbon (PGC) for a bed volume of 50 μL were washed with 500 μL of ddH2O, 500 μL of 80% acetonitrile (ACN), and equilibrated with 500 μL 0.1% trifluoroacetic acid (TFA). N-glycan pellets were dissolved in 50 μL of 0.1% TFA and loaded into the microtips at a flow rate of ~100 μL/min, washed with 500 μL 0.1% TFA, and N-glycans eluted with 500 μL of elution buffer (0.05% TFA, 40% ACN). The eluted N-glycans were analysed by LC-MS/MS. Total glycan samples were applied to a nano-HPLC Chip using a Agilent 1260 series microwell-plate autosampler, and interfaced with a Agilent 6550 iFunnel Q-TOF MS (Agilent Technologies, Inc., Santa Clara, CA). The HPLC Chip (glycan Chip) had a 40 nL enrichment column and a 75 μm x 43mm separation column packed with 5 μm graphitized carbon as stationary phase. The mobile phase was 0.1% formic acid in water (v/v) as solvent A, and 0.1% formic acid in ACN (v/v) as solvent B. The flow rate at 0.3 μL/min with gradient schedule; 5% B (0-1 min); 5-20% B (1-15 min); 20-70% B (15-16 min); 70% B (16-19 min) and 70-5% B (19-20 min). MS System was operated in positive ion mode at 2 GHz Extended Dynamic Range, MS mode in low mass range (1700 m/z) with MS setting at 8 MS (range 450–1700 m/z).

### MTT test

Thirty thousand cells dissociated were seeded on NFSs of various stiffnesses or in 2D. After 5 days of migration, 0.5μg/μl of MTT (Sigma) was added in the medium during 3 hours at 37°C/5%CO2. DMSO (sigma) was then added and the medium was transferred in 96 wells plates to be analysed by a Clariostar microplate reader.

### Cellular growth analysis

Ten thousand cells were seeded in 96 wells plates pre-coated with poly-2-hydroxyethyl methacrylate (poly-HEME, Sigma), and counted daily with Z2 counter (Beckman Coulter). Before counting, GSC were dissociated with Trypsin 2.5%/EDTA (10mM) and incubated 10min at 37°C/5%CO2 before particle counting.

### Immunostaining

After 5 days of culture, Gli4 on NFS or 2D were fixed by a solution of 4% of PFA. Cells were blocked and permeabilized using a solution of PBS - triton 0.5% - horse serum 5%. Primary antibodies were incubated overnight at 4°C. The antibody used in immunofluorescence was Vinculin (Sigma Aldrich V9264) Fluorochrome-coupled secondary antibodies (1/500) were incubated 2 hours at room temperature. The actin cytoskeleton was stained with phalloidin and cell nuclei with Hoechst 33342. NFS and coverslips with GSCs were mounted in fluoromount medium.

### Microscopy

Image capturing and Z-stack acquisition were performed using Confocal 2 Zeiss LSM 5 Live DUO and Widefield 1 – Zeiss Axioimager Z1/ Zen (with an apotome) microscopes. Imaris x64 8.1.2 software has been used for 3D image reconstitution. Quantifications were done using ZEN software. SEM images were performed using Hitachi S4800, Zeiss EVO HD15.

### Western Blot

Proteins were extracted by submerging the NFS in RIPA buffer (+ phosphatase/protease inhibitors). Protein lysate were separated by SDS-PAGE. PVDF membranes were blocked by TBS-Tween 0.1% - milk 5%. Primary antibodies were incubated overnight at 4°C. The antibodies used in western blot are: N-cadherin (N-CAD) (D4R1H) (Cell Signalling 13116S), Vimentin (D21H3) (Cell Signalling 5741S), ZO-1 (D7D12) (Cell Signalling 8193S), TCF8/ZEB1 (D80D3) (Cell Signalling 3396S), Calpain-2 (Abcam ab155666) Galectin-3 (Abcam ab2785), Talin1/2 (Abcam ab11188), FAK (Abcam ab40794), phospho-FAK Y397 (Abcam ab81298), SOX2 (Abcam ab97959), and GAPDH as a loading control (Millipore MAB374). Horseradish peroxidase-coupled secondary antibodies were incubated 2 hours at room temperature. The Chemidoc XRS+ imager was used for chemiluminescence detection. The pixel quantifications were done using Image Lab software.

### Statistical analysis

Forces were analysed using the Asylum Research software. All data are reported as mean ± standard error of the mean. ANOVA test was used for comparison between samples. Statistical analysis of the data was performed with Prism GraphPad.

## Acknowledgements

DC and NB thank INSERM for the financial support (INSERM/INCA project PC201216 Gliomatrack) and Jacques Vignon and Jan de Weille for the critical reading of the manuscript and the MENRT fellowship (CBS2 Doctoral School).

---

**Supplementary figure 1:**
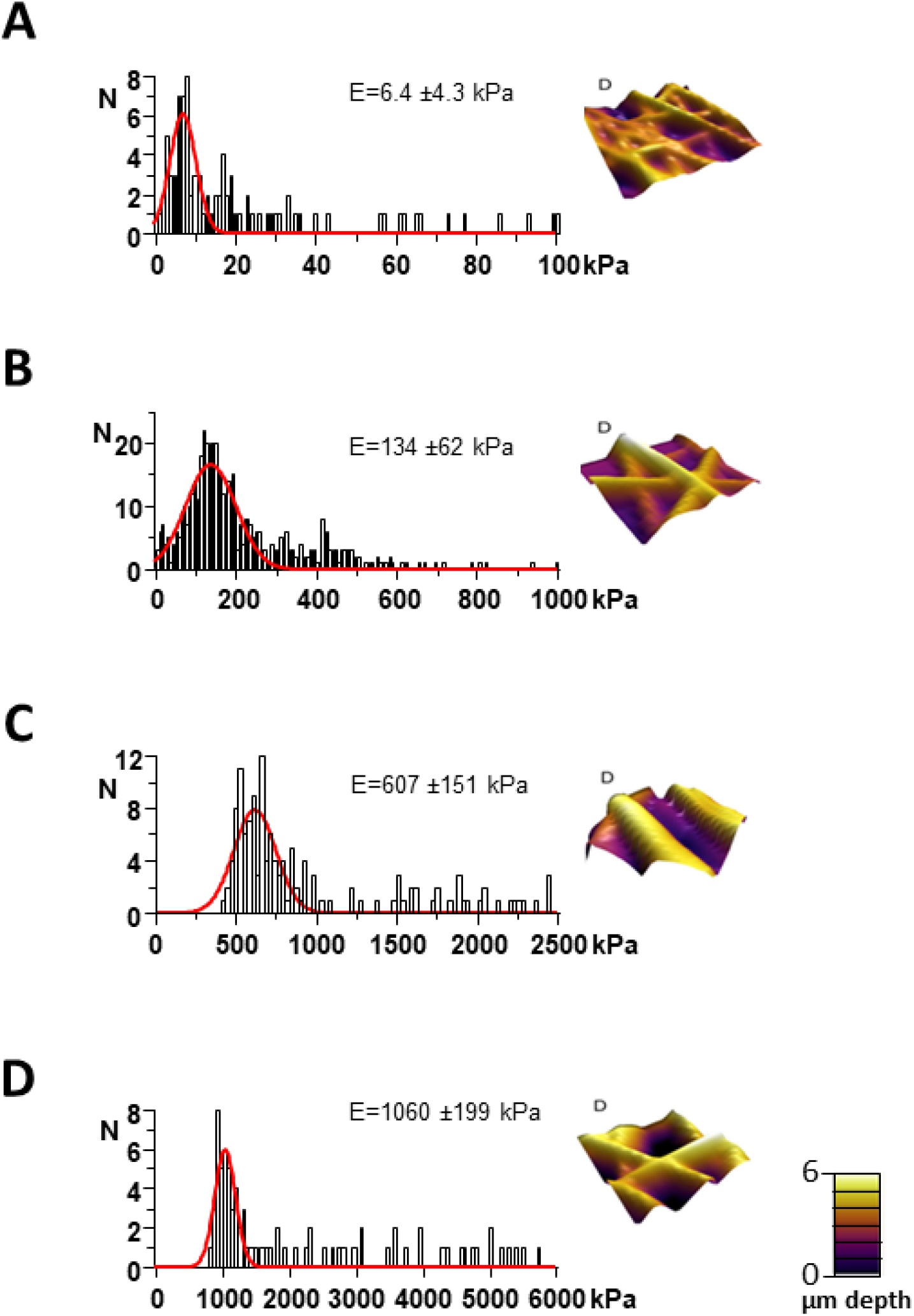
Atomic Force Microscopy force-volume maps. (A, B, C, D) 3D reconstructions of height of the contact point of control, 0.0015% w/w of MWCNT, 0.00625% w/w and 0.05% w/w samples, respectively. Histograms of the associated Young’s modulus E fitted with a Gaussian distribution. Stiffness values given in the text correspond to the mean ± SD of 5 independent elasticity maps. Statistical significance was determined using one-way ANOVA with post hoc Tukey’s Honest Significant Difference test for multiple comparisons. P values of less than 0.05 were considered significant. (D) 3D-reconstruction of the contact point’s height.

**Supplementary figure 2:**
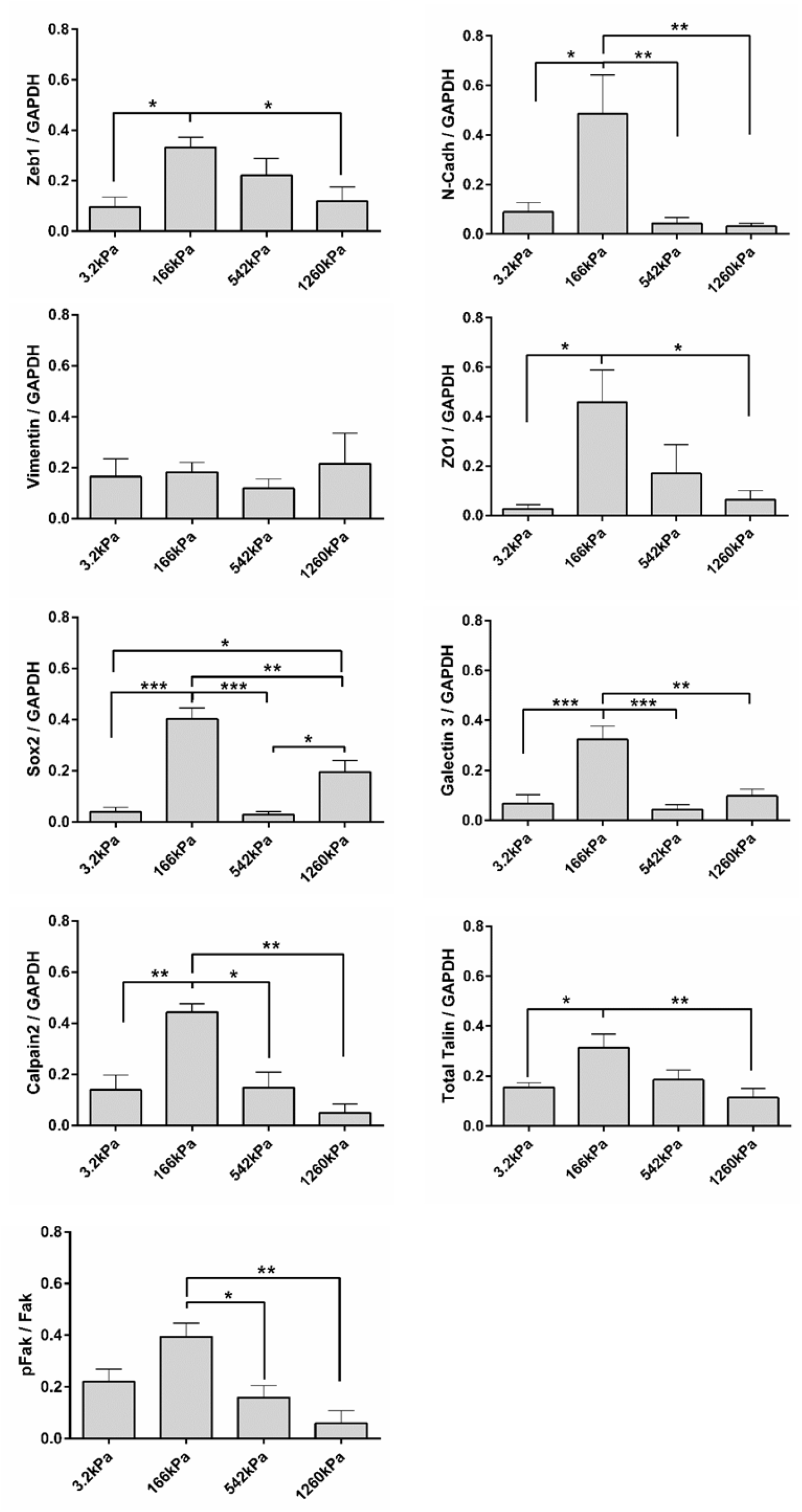
Western Blot quantification of WT GSC proteins expression. Each western blot quantification was performed three to seven times and normalized with respect to GAPDH expressions and sum. Bands were quantified using the Bio-Rad Chemidoc and the image lab software. Data are presented as mean +/- SEM and statistical significance was determined using one-way ANOVA (* p<0,05; ** p<0,01; *** p<0,005; **** p<0,001).

**Supplementary figure 3:**
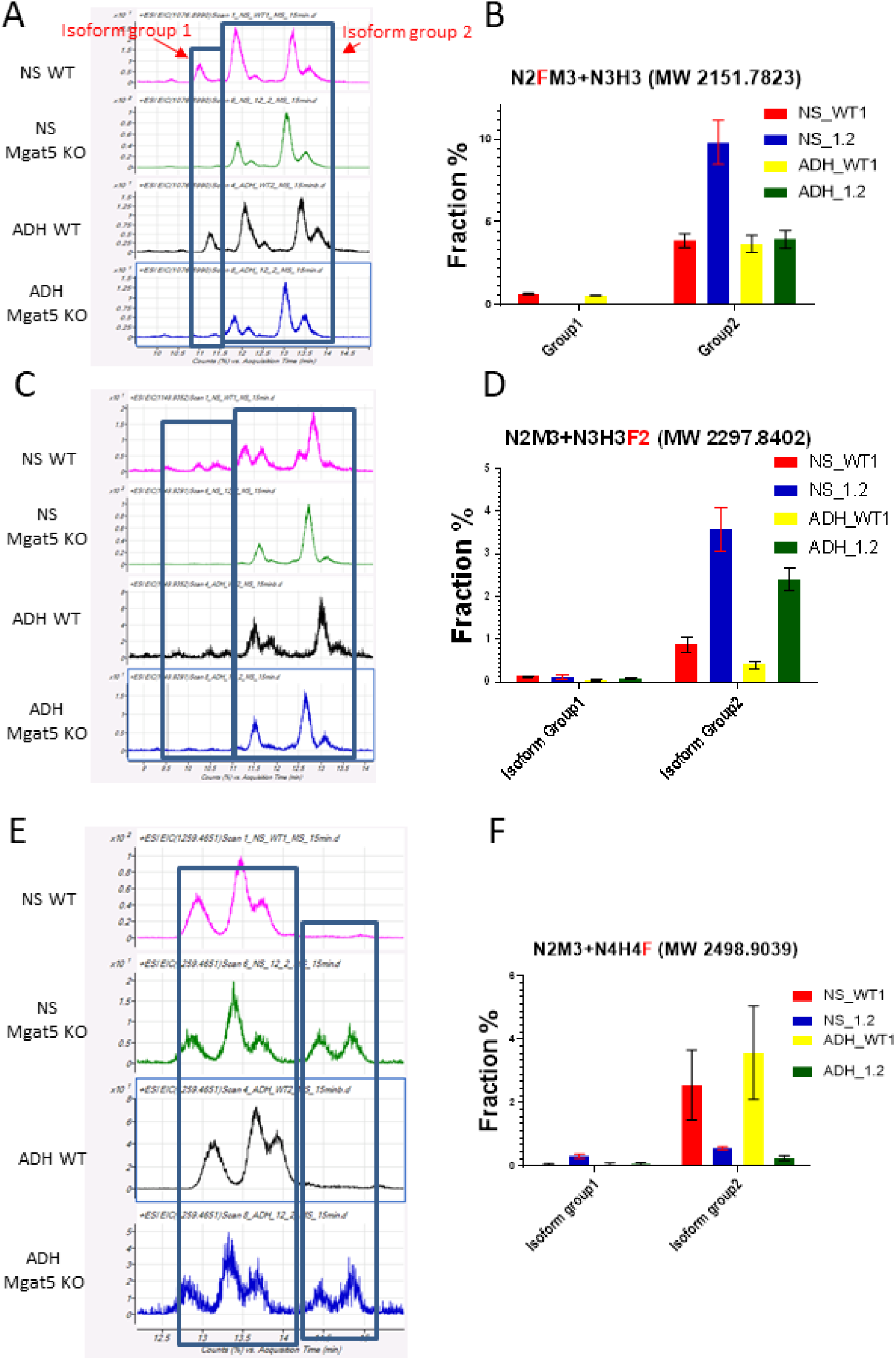
Examples of MS analysis and quantification of fucosylated glycans. Glycans were isolated from wild type (WT) or MGAT5 KO (1.2) GSCs cultivated either as neurospheres (NS) in proliferation medium or as adherent cells (ADH) in 2D in differentiation medium. The minor structures with identical masses, group 2 and 1 are likely to be isomers, for example two linear N-acetyllactosamine units on a bi- rather than tri – antennary N-glycan. (A, B) MS analysis and quantification tri-antenate glycancs with a fucosylated core of. (B, C, D) MS analysis and quantification of tri-antenate glycans di-fucosylated on the branches. (E, F) MS analysis and quantification tetra-antenate glycans mono fucosylated on the branches. Data are presented as mean +/- SEM

**Supplementary figure 4:**
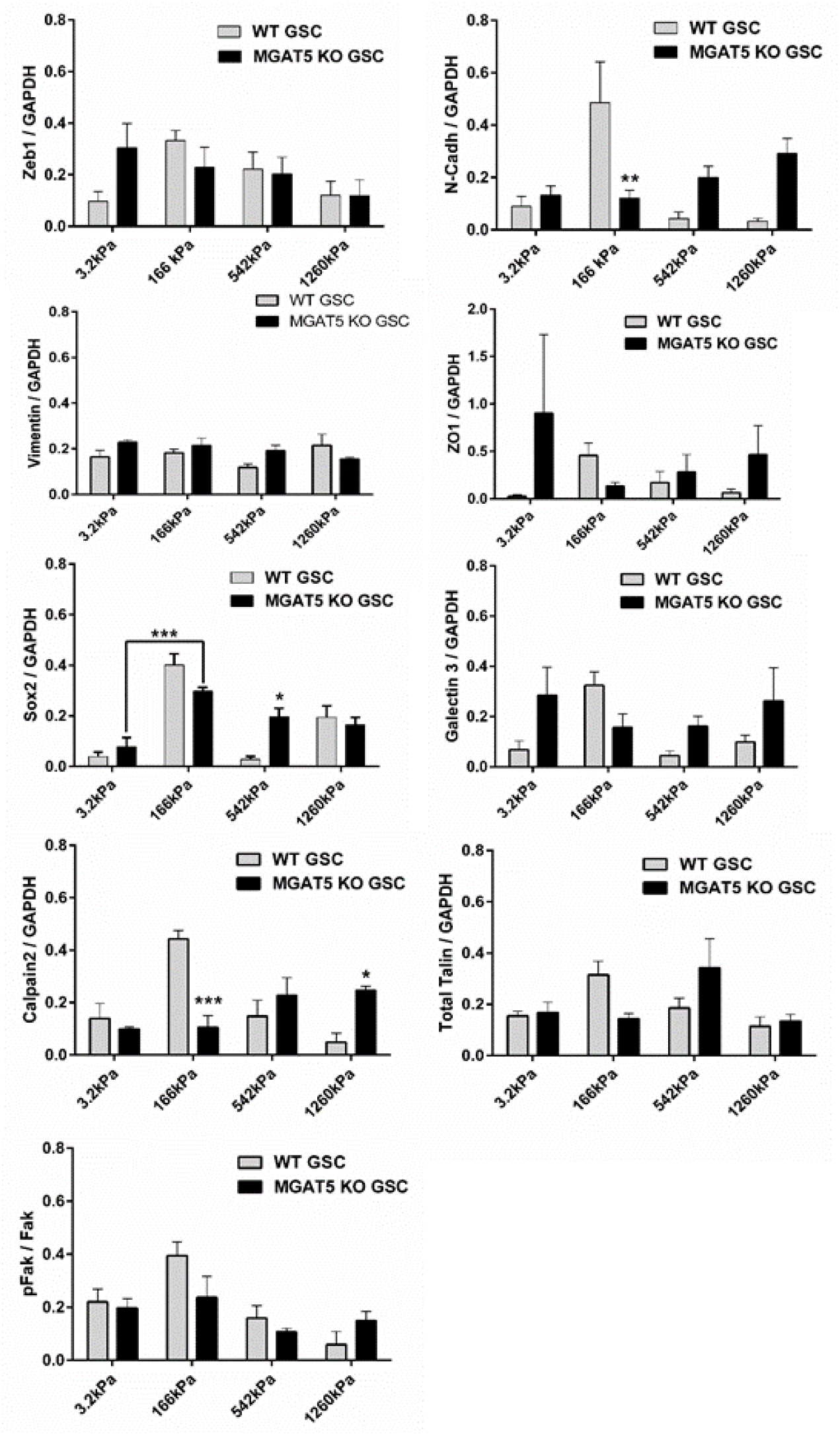
Western Blot quantification of MGAT5 KO GSC proteins expression. Each western blot quantification was performed three to four times and normalized with respect to GAPDH expressions and sum. Bands were quantified using the Bio-Rad Chemidoc and the image lab software. Data are presented as mean +/- SEM and statistical significance was determined using one-way ANOVA (* p<0,05; ** p<0,01; *** p<0,005; **** p<0,001).

**Supplementary figure 5:**
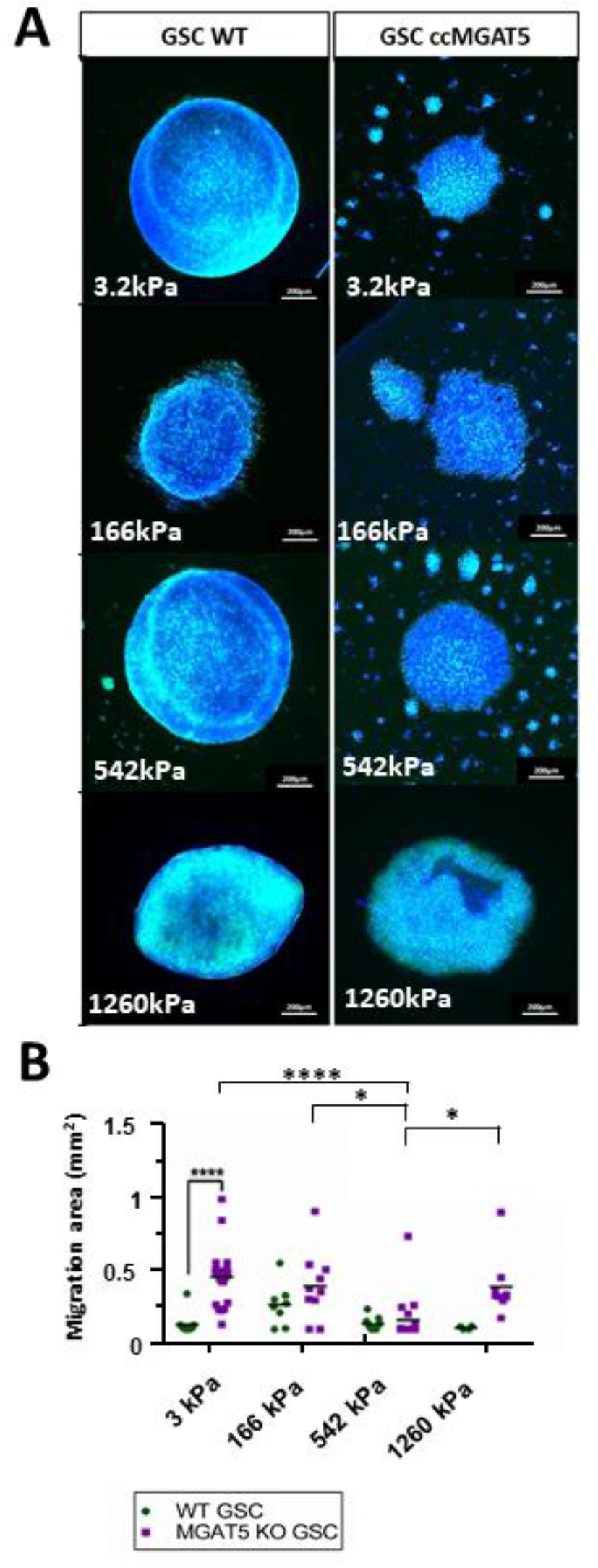
MGAT5 KO GSC migration in proliferation medium compared to WT GSCs. (A) Pictures of WT and MGAT5 KO WT GSCs migration behaviour in proliferation medium of WT GSCs at every stiffnesses. Nucleus stain with Hoechst 33342. (B) Quantification 5 days after NS plating of migration area (μm^2^) in the proliferation condition of WT GSCs and MGAT5 KO GSCs. Data are presented as mean +/- SEM and statistical significance was determined using one-way ANOVA (* p<0,05; ** p<0,01; *** p<0,005; **** p<0,001).

## References

1. Y. L. Hu et al., FAK and paxillin dynamics at focal adhesions in the protrusions of migrating cells. Sci Rep 4, 6024 (2014).

2. E. R. Horton et al., The integrin adhesome network at a glance. J Cell Sci 129, 4159–4163 (2016).

3. T. Mitchison, M. Kirschner, Cytoskeletal dynamics and nerve growth. Neuron 1, 761–772 (1988).

4. A. Elosegui-Artola et al., Rigidity sensing and adaptation through regulation of integrin types. Nat Mater 13, 631–637 (2014).

5. B. T. Goult, J. Yan, M. A. Schwartz, Talin as a mechanosensitive signaling hub. J Cell Biol 217, 3776–3784 (2018).

6. A. Carisey et al., Vinculin regulates the recruitment and release of core focal adhesion proteins in a force-dependent manner. Curr. Biol. 23, 271–281 (2013).

7. M. S. Bauer et al., Structural and mechanistic insights into mechanoactivation of focal adhesion kinase. Proc. Natl. Acad. Sci. U. S. A. 116, 6766–6774 (2019).

8. P. C. Kerstein, K. M. Patel, T. M. Gomez, Calpain-Mediated Proteolysis of Talin and FAK Regulates Adhesion Dynamics Necessary for Axon Guidance. J Neurosci 37, 1568–1580 (2017).

9. J. G. Goetz et al., Concerted regulation of focal adhesion dynamics by galectin-3 and tyrosine-phosphorylated caveolin-1. J Cell Biol 180, 1261–1275 (2008).

10. K. S. Lau et al., Complex N-glycan number and degree of branching cooperate to regulate cell proliferation and differentiation. Cell 129, 123–134 (2007).

11. E. A. Partridge et al., Regulation of cytokine receptors by Golgi N-glycan processing and endocytosis. Science 306, 120–124 (2004).

12. A. Lagana et al., Galectin binding to Mgat5-modified N-glycans regulates fibronectin matrix remodeling in tumor cells. Mol Cell Biol 26, 3181–3193 (2006).

13. J. M. Barnes, L. Przybyla, V. M. Weaver, Tissue mechanics regulate brain development, homeostasis and disease. J Cell Sci 130, 71–82 (2017).

14. V. A. Cuddapah, S. Robel, S. Watkins, H. Sontheimer, A neurocentric perspective on glioma invasion. Nat Rev Neurosci 15, 455–465 (2014).

15. S. Watkins, H. Sontheimer, Hydrodynamic cellular volume changes enable glioma cell invasion. J Neurosci 31, 17250–17259 (2011).

16. H. J. Scherer, Structural development in gliomas. Am J Cancer 34, 333–351 (1938).

17. R. Malik, P. I. Lelkes, E. Cukierman, Biomechanical and biochemical remodeling of stromal extracellular matrix in cancer. Trends Biotechnol 33, 230–236 (2015).

18. A. Saleh et al., A novel 3D nanofibre scaffold conserves the plasticity of glioblastoma stem cell invasion by regulating galectin-3 and integrin-beta1 expression. Sci Rep 9, 14612 (2019).

19. D. C. Stewart, A. Rubiano, K. Dyson, C. S. Simmons, Mechanical characterization of human brain tumors from patients and comparison to potential surgical phantoms. PLoS One 12, e0177561 (2017).

20. M. C. Weisenberger, E. A. Grulke, D. Jacques, T. Rantell, Enhanced mechanical properties of polyacrilonitrile/multiwall carbon nanotube composites fibers. J. Nanosci. Nanotechnol. 3, 535–539 (2003).

21. F. A. Siebzehnrubl, B. A. Reynolds, A. Vescovi, D. A. Steindler, L. P. Deleyrolle, The origins of glioma: E Pluribus Unum? Glia 59, 1135–1147 (2011).

22. C. Beadle et al., The role of myosin II in glioma invasion of the brain. Mol Biol Cell 19, 3357–3368 (2008).

23. R. J. Petrie, A. D. Doyle, K. M. Yamada, Random versus directionally persistent cell migration. Nat Rev Mol Cell Biol 10, 538–549 (2009).

24. W. Xu et al., Cell stiffness is a biomarker of the metastatic potential of ovarian cancer cells. PLoS One 7, e46609 (2012).

25. M. J. Paszek et al., The cancer glycocalyx mechanically primes integrin-mediated growth and survival. Nature 511, 319–325 (2014).

26. V. Swaminathan, R. S. Fischer, C. M. Waterman, The FAK-Arp2/3 interaction promotes leading edge advance and haptosensing by coupling nascent adhesions to lamellipodia actin. Mol Biol Cell 27, 1085–1100 (2016).

27. J. G. Goetz, Bidirectional control of the inner dynamics of focal adhesions promotes cell migration. Cell Adh Migr 3, 185–190 (2009).

28. T. Doetschman, T. Georgieva, Gene Editing With CRISPR/Cas9 RNA-Directed Nuclease. Circ Res 120, 876–894 (2017).

29. Z. Hassani et al., Phostine PST3.1a Targets MGAT5 and Inhibits Glioblastoma-Initiating Cell Invasiveness and Proliferation. Mol Cancer Res 15, 1376–1387 (2017).

30. S. S. Pinho, C. A. Reis, Glycosylation in cancer: mechanisms and clinical implications. Nat Rev Cancer 15, 540–555 (2015).

31. K. S. Lau, J. W. Dennis, N-Glycans in cancer progression. Glycobiology 18, 750–760 (2008).

32. H. Guo et al., Transcriptional regulation of the protocadherin beta cluster during Her-2 protein-induced mammary tumorigenesis results from altered N-glycan branching. J Biol Chem 287, 24941–24954 (2012).

33. M. Saxena, R. Changede, J. Hone, H. Wolfenson, M. P. Sheetz, Force-Induced Calpain Cleavage of Talin Is Critical for Growth, Adhesion Development, and Rigidity Sensing. Nano Lett 17, 7242–7251 (2017).

34. M. Bosch-Fortea, F. Martin-Belmonte, Mechanosensitive adhesion complexes in epithelial architecture and cancer onset. Curr Opin Cell Biol 50, 42–49 (2018).

35. B. Klapholz, N. H. Brown, Talin - the master of integrin adhesions. J Cell Sci 130, 2435–2446 (2017).

36. A. Huttenlocher, A. R. Horwitz, Integrins in cell migration. Cold Spring Harb Perspect Biol 3, a005074 (2011).

37. D. S. Harburger, D. A. Calderwood, Integrin signalling at a glance. J Cell Sci 122, 159–163 (2009).

38. D. Ilic et al., Reduced cell motility and enhanced focal adhesion contact formation in cells from FAK-deficient mice. Nature 377, 539–544 (1995).

39. P. Euskirchen et al., Cellular heterogeneity contributes to subtype-specific expression of ZEB1 in human glioblastoma. PLoS One 12, e0185376 (2017).

40. Y. Chen et al., ZEB1 Regulates Multiple Oncogenic Components Involved in Uveal Melanoma Progression. Sci Rep 7, 45 (2017).

41. P. Friedl, R. Mayor, Tuning Collective Cell Migration by Cell-Cell Junction Regulation. Cold Spring Harb Perspect Biol 9 (2017).

42. W. Zhou et al., Cancer-secreted miR-105 destroys vascular endothelial barriers to promote metastasis. Cancer Cell 25, 501–515 (2014).

43. L. Chen et al., Metastasis is regulated via microRNA-200/ZEB1 axis control of tumour cell PD-L1 expression and intratumoral immunosuppression. Nat Commun 5, 5241 (2014).

44. T. Svaldo Lanero, O. Cavalleri, S. Krol, R. Rolandi, A. Gliozzi, Mechanical properties of single living cells encapsulated in polyelectrolyte matrixes. J Biotechnol 124, 723–731 (2006).

45. H. Hertz, Ueber die Berührung fester elastischer Körper. Journal für die reine und angewandte Mathematik 92, 156–171 (1881).

46. M. Martin et al., Morphology and nanomechanics of sensory neurons growth cones following peripheral nerve injury. PLoS One 8, e56286 (2013).

47. M. Radmacher, M. Fritz, C. M. Kacher, J. P. Cleveland, P. K. Hansma, Measuring the viscoelastic properties of human platelets with the atomic force microscope. Biophys J 70, 556–567 (1996).

48. T. Boudou, J. Ohayon, C. Picart, P. Tracqui, An extended relationship for the characterization of Young’s modulus and Poisson’s ratio of tunable polyacrylamide gels. Biorheology 43, 721–728 (2006).

49. P. O. Guichet et al., Cell death and neuronal differentiation of glioblastoma stem-like cells induced by neurogenic transcription factors. Glia 61, 225–239 (2013).

50. D. K. Singh et al., Oncogenes Activate an Autonomous Transcriptional Regulatory Circuit That Drives Glioblastoma. Cell Rep 18, 961–976 (2017).

